# Spatial Clustering of Interface Residues Enhances Few-Shot Prediction of Viral Protein Binding

**DOI:** 10.1101/2025.04.10.647895

**Authors:** Maxime Basse, Dianzhuo Wang, Eugene I. Shakhnovich

## Abstract

Predicting protein binding affinities across large combinatorial mutation spaces remains a critical challenge in molecular biology, particularly for understanding viral evolution and antibody interactions. While combinatorial mutagenesis experiments provide valuable data for training predictive models, they are typically limited due to experimental constraints. This creates a significant gap in our ability to predict the effects of more extensive mutation combinations, such as those observed in emerging SARS-CoV-2 variants. We present PROXICLUST, which strategically combines smaller combinatorial mutagenesis experiments to enable accurate predictions across larger combinatorial spaces. Our approach leverages the spatial proximity of amino acid residues to identify potential epistatic interactions, using these relationships to optimize the design of manageable-sized combinatorial experiments. By combining just two small combinatorial datasets, we achieve accurate binding affinity predictions across substantially larger mutation spaces (*R*^2^ ≈ 0.8), with performance strongly correlated with capture of high-order epistatic effects. We validated our method in five different protein-protein interaction datasets, including binding of SARS-CoV-2 receptor binding domain (RBD) to various antibodies and cellular receptors, as well as influenza RBD-antibody interactions. This work provides a practical framework for extending the predictive power of combinatorial mutagenesis beyond current experimental constraints, offering applications in viral surveillance and antibody engineering.

## 1 Introduction

Predicting the dissociation constant (*K*_*D*_) between viral proteins and monoclonal antibodies, as well as the cellular receptor Angiotensin-converting enzyme 2 (ACE2), is critical for assessing viral infectivity for viruses like SARS-CoV-2, as demonstrated in these works Wang et al. (2024); Cia et al. (2022); Hie et al. (2021); Huot et al. (2025). These models, whether based on biophysics or machine learning, utilize the *K*_*D*_ of various antibodies and the receptor-binding domain (RBD) to effectively predict viral fitness.

High-throughput experimental methods provide a means to acquire *K*_*D*_ data across extensive sequence spaces. A prominent technique is Combinatorial Mutagenesis Moulana et al. (2023; 2022). In these experiments, researchers select *L* specific loci within the Receptor Binding Domain (RBD), testing every possible combination of amino acid mutations at each locus. This method generates a comprehensive dataset comprising 2^*L*^ combinations, capturing a broad spectrum of mutational impacts. This methodology is particularly crucial in the context of viral evolution, where early in a pandemic, variants of concern (VoCs) with specific mutations are observed Deng et al. (2021); Leung et al. (2021). It is vital to predict the phenotypic or fitness outcomes of variants combining these mutations, as illustrated in Figure 1. However, currently, the capabilities of these methods are restricted to experiments where *L* ≤ 16, due to the exponential growth in the number of combinations with larger values of *L*. VoCs such as BA.2, BA.3 and XBB variants Tamura et al. (2024), often exhibit more than 16 mutations Carabelli et al. (2023), underscoring a gap between current experimental capabilities and the need to effectively predict infectivity of VOCs with more mutations.

**Figure 1:**
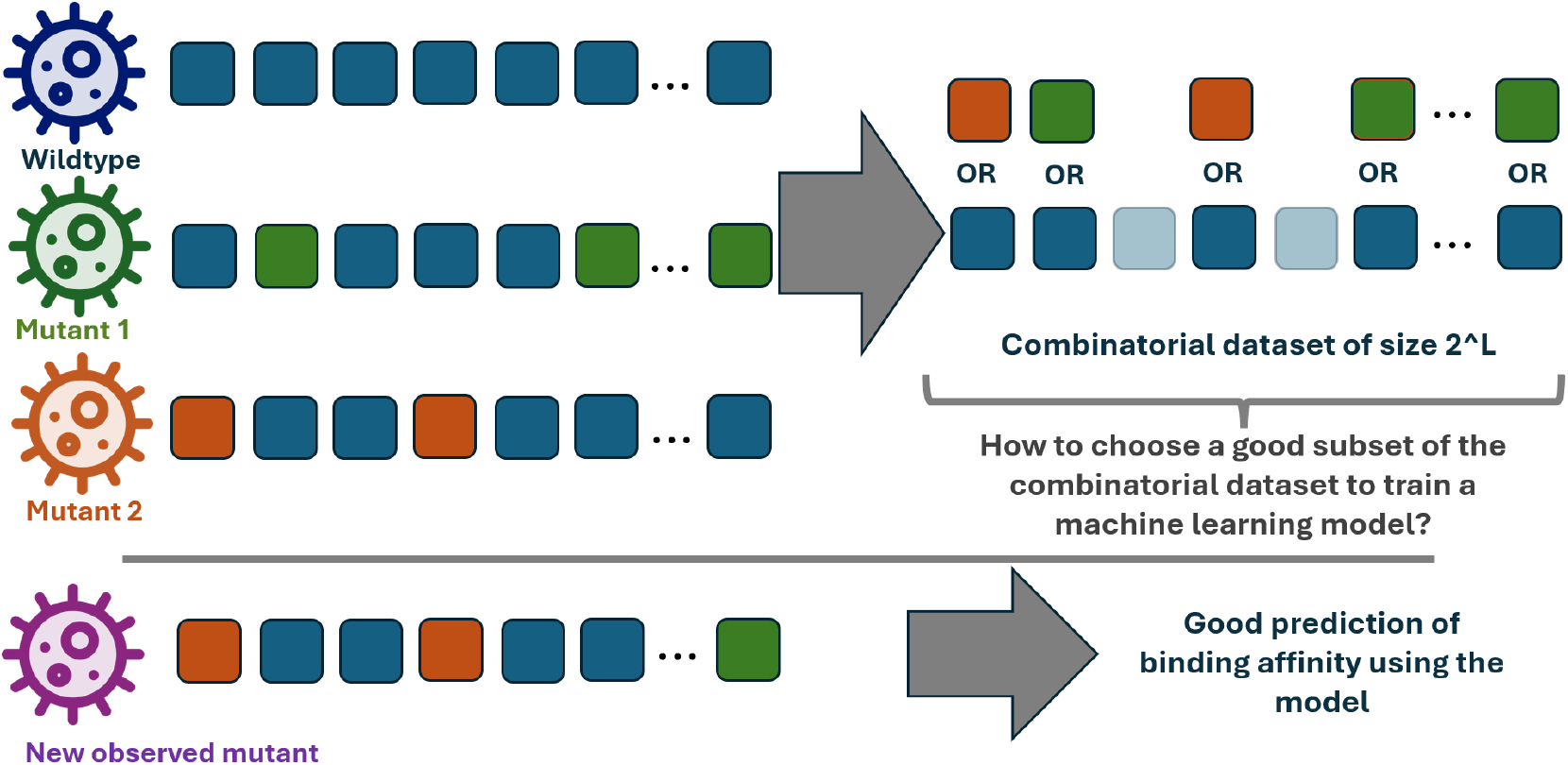
A more infectious Variant of Concern (VoC) may incorporate mutations from existing VoCs. Consequently, it is crucial to predict the biophysical properties of the entire combination of existing VoCs to better understand and anticipate their evolutionary trajectories.

Figure 1 illustrates how a combinatorial set of Variants of Concern (VOCs) can be derived from mutations observed in circulating viral strains. Accurate predictions across this combinatorial space are essential for reconstructing viral fitness landscapes, as these sets encompass all possible intermediary variants between wildtype and observed mutants. The mutations included in these sets are particularly significant because they originate from viruses that have demonstrated successful transmission and survival in natural populations. Consequently, new variants combining these individually beneficial mutations have an increased likelihood of being both infectious and viable. While the combinatorial space of 2^*L*^ variants represents only a small fraction of the total possible sequence space (20^*K*^, where *K* is the total number of residues in the viral protein), it is enriched for potentially concerning variants due to this selective sampling of successful mutations. The ability to accurately predict *K*_*D*_ and fitness across this space is therefore crucial for proactive vaccine development, therapeutic adaptation, and evidence-based public health policy decisions.

During the early stages of a pandemic, when experimental capacity is often limited relative to the urgent need for data, traditional experimental methods alone may not be sufficient to characterize all emerging viral variants. In these scenarios, machine learning approaches trained on limited available experimental data have proven to be valuable tools for estimating *K*_*D*_ of novel variantsWang et al. (2024); Han et al. (2023); Loux et al. (2024). This computational approach helps bridge the gap between experimental capabilities and the pressing need for rapid viral surveillance.

Significant advances have been made in computational methods for predicting *K*_*D*_ to address limitations in experimental throughput. These approaches span a broad methodological spectrum, from transformer-based architectures Wang et al. (2023) and physics-informed deep learning models Chen et al. (2021) to free energy perturbation calculations Sergeeva et al. (2023). Complementing these modeling advances, active learning (AL) strategies have emerged as powerful tools for optimizing experimental data collection. AL enable strategic sampling from the vast space of possible variants, prioritizing measurements that maximize information gain for model improvement. This could significantly reduce the amount of training data required while maintaining high model accuracy Hie et al. (2020). However, a key limitation of active learning is its inherently sequential nature - experiments must be conducted one after another based on model feedback. This sequential constraint makes AL incompatible with high-throughput experimental methods that require upfront design of mutation libraries. The benchmark of AL and other low-throughput sampling methods could be found in SI.

To overcome this fundamental limitation, we investigate optimal methods for combining smaller, experimentally feasible combinatorial datasets to enable accurate predictions across larger combinatorial spaces. To this end, we introduce “PROXICLUST”, which leverages structural information from protein-protein binding poses to systematically determine which mutations should be grouped together in combinatorial experiments.

## 2 Methods

### PROXICLUST

We introduce PROXICLUST, a method based on the principle that combinatorial experiments reveal high-order epistatic interactions among mutations. Our key insight is that mutations with potential interactions should be tested together in experimental designs. These interacting mutations typically form “epistasis clusters” - groups of residues that functionally influence each other. While precise epistatic maps of viruses are rarely available, we leverage the observation that spatial proximity between amino acids serves as a reliable predictor of epistatic interactions Wang et al. (2024). PROXICLUST requires only structural information from the antibody-antigen binding pose, typically available through Protein Data bank Format files of wildtype variants, and proceeds in two steps:

Our method proceeds in two steps. First, we identify interface residues between the antibody-antigen complex by analyzing spatial coordinates in the binding pose. These interface residues can be determined either computationally using protein structure prediction methods like AlphaFold Multimer Jumper et al. (2021), or experimentally through cryo-electron microscopy. As in standard protein-protein interaction studies, we define interface residues as those within a 10Å distance threshold of the binding partner.

Second, we apply a two-stage clustering approach to these interface residues (Figure 2b). We first employ M-means clustering to establish well-distributed centroids across the binding interface, followed by an L-Nearest Neighbors algorithm initiated from these centroids. This hierarchical approach, implemented using scikit-learn’s KMeans and KNeighborsClassifier Pedregosa et al. (2011), creates partially overlapping clusters that capture local structural relationships while maintaining diversity across the interface. The overlap between clusters is carefully controlled through centroid initialization to maximize coverage of potential epistatic interactions while minimizing redundant measurements, as validated by our epistasis analysis (Figure 4).

**Figure 2:**
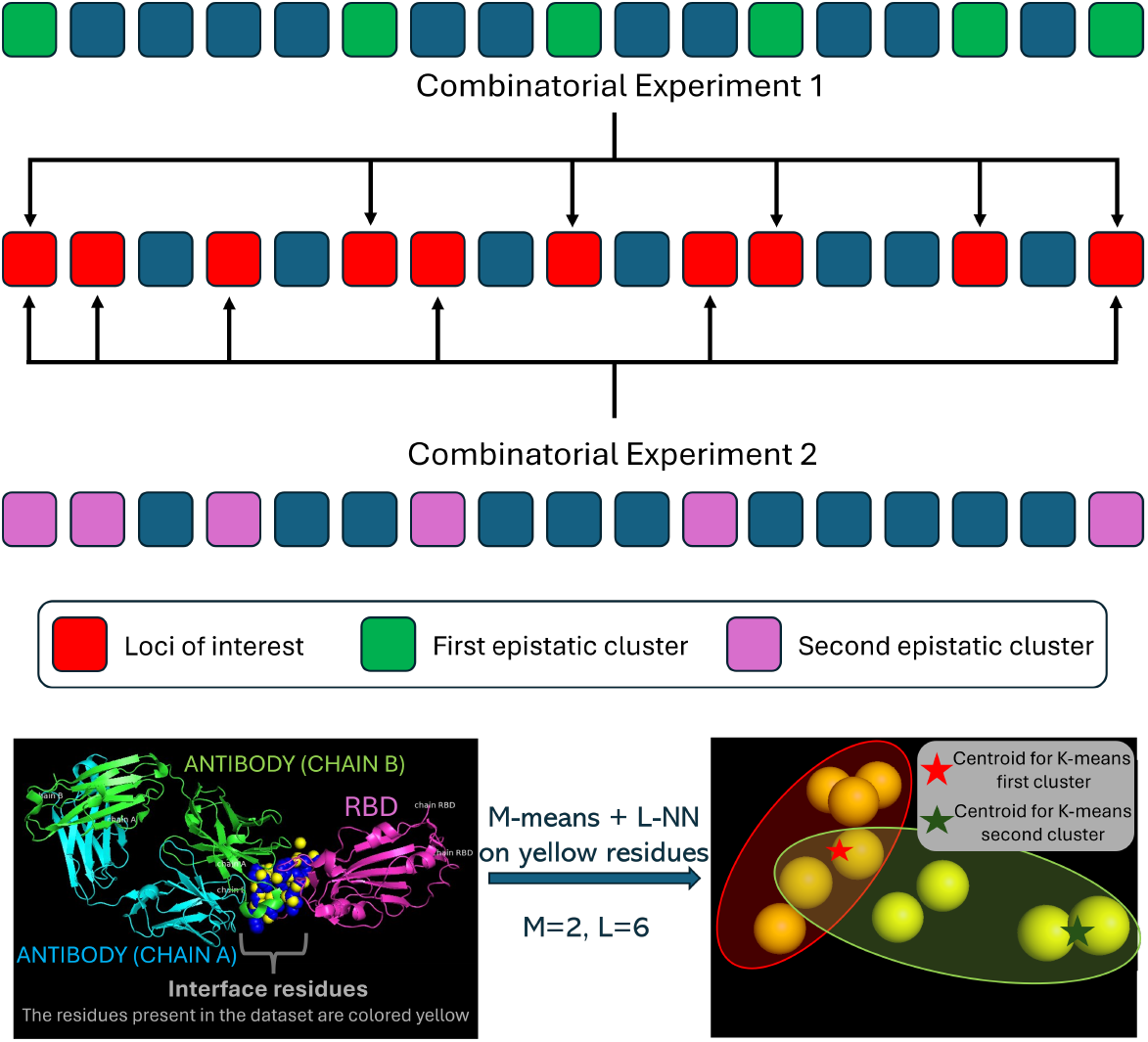
(a) Illustration of using two smaller combinatorial experiments to predict the outcomes of a larger experiment that is out of current capabilities (b) PROXICLUST employs M-means clustering and L-nearest neighbors algorithms to determine which amino acids should be grouped together in the same experiment.

The algorithm ultimately organizes interface residues into M groups of L residues each, where L represents the maximum number of mutations that can be included in a single combinatorial experiment (determined by experimental constraints), and M is the desired number of parallel experiments. Throughout this study, we demonstrate results for *M* = 2, showing that just two well-designed combinatorial experiments can effectively capture the key epistatic interactions in the system.

### DATASETS

In this study, we analyzed five datasets, including the combinatorial datasets of SARS-CoV2 RBD binding with ACE2 Moulana et al. (2022), LY-CoV016, LY-CoV555, REGN10987, and S309 Moulana et al. (2023), as well as the anti-influenza receptor binding site (RBS), CH65 binding to H1 Phillips et al. (2023).

The SARS-CoV-2 datasets Moulana et al. (2022; 2023) explore the combinations of 15 mutations on the RBD corresponding to the 15 mutations observed on Omicron BA.1 variant compared to the Wuhan wildtype: G339D, S371L, S373P, S375F, K417N, N440K, G446S, S477N, T478K, E484A, Q493R, G496S, Q498R, N501Y, Y505H. The influenza dataset Phillips et al. (2023) contains all possible evolutionary intermediates between the unmutated common ancestor and the mature somatic sequence of the CH65 antibody, which binds to diverse H1 strains. The dataset contains all combinations of 16 mutations located on both the heavy and light chain of the CH65 antibody.

These datasets are comprehensive combinatorial mutagenesis scans that measure binding affinity values for all possible combinations of selected mutations. Each mutant variant is represented using a binary one-hot encoding scheme, where ‘1’ indicates the presence of a mutation at a specific position and ‘0’ represents the wild-type residue, as illustrated in Table 1. For a set of L selected mutation sites, this generates a complete combinatorial space of 2^*L*^ variants, systematically exploring all possible mutational combinations.

## 3 Results

In this section, we address the challenge of assembling multiple smaller combinatorial scans into a training set to predict a larger combinatorial dataset. We benchmark PROXICLUST against random assembly strategies of the same dimensionality. Here, a random assembly strategy refers to selecting mutations for each scan independently as a random subset from the pool of possible mutations.

Figure 3 compares the performance of a random forest machine learning model using two distinct strategies for assembling combinatorial datasets:

**Figure 3:**
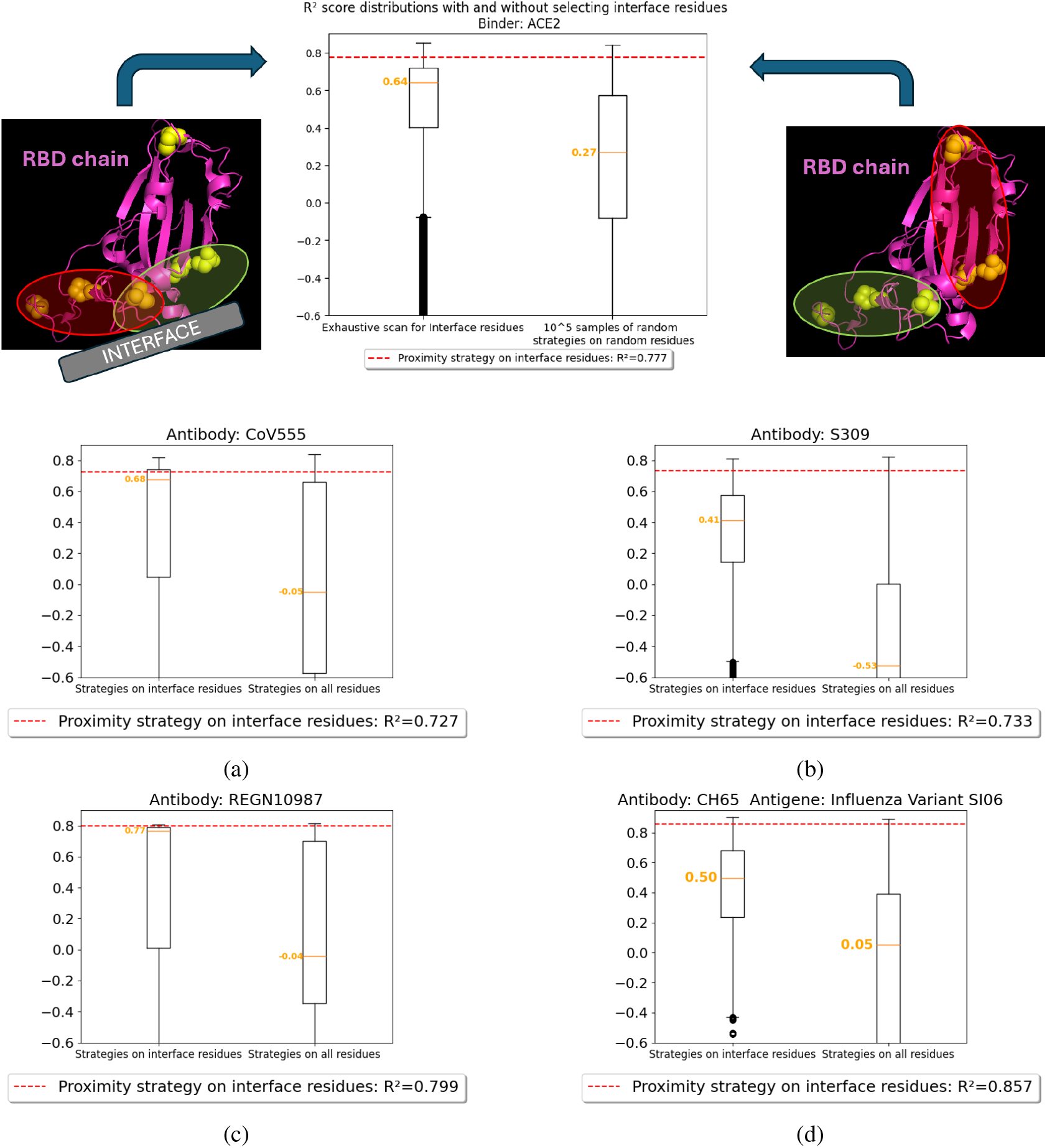
Comparison of *R*^2^ scores of models trained using data acquisition strategies focused on the antibody-antigen interface (left boxplots) versus unconstrained strategies (right boxplots). Red line shows the *R*^2^ score achieved by PROXICLUST. (a) Comparison for RBD and ACE2 binding. The accompanying protein illustrations represent possible clustering of the dataset’s residues shown with yellow spheres. On the left, the clustering is done among the yellow spheres close to the interface whereas on the right, all residues can be included in the clusters. (b)-(d) Similar analyses were conducted for RBD binding with antibodies LY-CoV555, S309, and REGN10987, respectively. (e) Same experiment run on another combinatorial dataset for binding affinity predictions between Influenza variant SI06 and variants of antibody CH65

- The boxplots on the left display the distribution of *R*^2^ scores for strategies that assemble two combinatorial scans, each targeting mutations specifically at the interface of the antigen-antibody binding pose.
- The boxplots on the right show the distribution of *R*^2^ scores for strategies that assemble two combinatorial scans, but with no constraints on the mutations included.

Comparing the medians in Figure 3, indicated by the orange horizontal lines, we observe that strategies targeting the interface consistently achieve higher scores. This finding aligns with our current understanding of protein-protein interactions - residues at binding interfaces are known to make the most significant contributions to binding affinity through direct physical contacts Moulana et al. (2022); Vangone & Bonvin (2015). The superior performance of interface-focused strategies can thus be attributed to their ability to capture these critical first-order effects in the training data. By concentrating on interface residues, these approaches naturally prioritize the most functionally relevant mutations, leading to more accurate predictions of binding affinity changes.

Our analysis reveals that PROXICLUST consistently outperforms both the median combinatorial scan assembly and the median interface assembly strategy (Figure 3). A detailed performance breakdown across all five tested datasets is presented in SI Table 2, where quantile rankings demonstrate how our method compares against a comprehensive pool of randomized residue selection strategies. Further detailed comparisons within interface assembly strategies are visualized in SI Figure 6. The performance advantage of PROXICLUST varies notably across different systems. For REGN10987 and LY-CoV555 antibodies, where the *R*^2^ score distributions peak around 0.8, the improvement over median performance is modest (*≤*0.3). However, for the Influenza virus and S309 antibody systems, PROXICLUST demonstrates a more substantial advantage, achieving *R*^2^ scores of approximately 0.8 compared to random strategies’ scores of around 0.4. This variable performance across different systems suggests that the effectiveness of our method may depend on the underlying structural and biochemical properties of the protein-protein interaction being studied.

Note that the assembling strategies being compared are always of the same “dimension”, meaning they consist of an equal number of combinatorial scans, each of the same size. This uniformity is crucial as, aside from the assembly strategy, the size of the training set is a key determinant in the performance of the predictive model. Therefore, it is essential to compare strategies that provide training sets of comparable sizes to ensure valid conclusions about their efficacy.

It is important to note that while strategies of the same dimension incur identical experimental costs (requiring the same number and type of laboratory experiments), they can yield different numbers of unique binding affinity measurements. To illustrate this concept, consider two contrasting scenarios: In the first case, a strategy using two scans with completely distinct sets of 5 mutations each generates 2^5^ + 2^5^ = 64 unique measurements. In contrast, a strategy of identical dimension but with 4 overlapping mutations between scans yields fewer unique measurements - 2^5^ measurements from the first scan plus only 2^5^*−* 2^4^ = 16 additional unique measurements from the second scan, totaling 48 unique mutants in the training set. This difference in measurement efficiency arises from the redundancy introduced by overlapping mutations, despite equivalent experimental effort.

### EPISTASIS ANALYSIS

We have developed a method to quantify the epistasis captured by a training set derived from any given assembly strategy. This method leverages the Walsh-Hadamard transform.

Walsh-Hadamard transform Poelwijk et al. (2016) is a method adapted from signal processing to study high-order epistasis(non-linearity) Faure et al. (2024). This approach is particularly advantageous when working with complete combinatorial datasets, as it provides exhaustive information about the interactions between residues Weinreich et al. (2013). Given that our binding affinity datasets are combinatorial in nature (of size 2^*L*^), we can derive the effect of any order of epistasis by applying the Walsh-Hadamard transform to the vector of experimentally measured binding affinities.

In the matrix form, this could be written as:

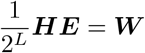

where H is the Hadamard matrix with size 2^*L*^ *×* 2^*L*^, and E is the Kd vector with size 2^*L*^*×* 1. W is the Walsh coefficients vector with size 2^*L*^ *×* 1, where each component *W*_*g*_ represents the importance (or weight) of a specific genotype *g*Weinreich et al. (2013).

A data acquisition strategy composed of multiple combinatorial scans can be presented as [*l*_1_, …, *l*_*M*_], representing the assembly of *M* individual combinatorial scans. Each *l*_*k*_ is a subset of {1, …, *L*} that specifies the positions (or loci) in a genotype where mutations (represented by 1s) can occur. A genotype *g* in {0, 1}^*L*^ is a sequence of binary values, where a 1 indicates the presence of a mutation at a given position, and a 0 indicates no mutation.

We define *G*_*k*_ as the set of all genotypes *g* in the combinatorial mutagenesis scan on the loci of *l*_*k*_. Formally:

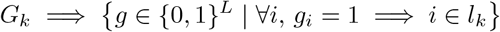

#### Example

Suppose *L* = 5 and *l*_*k*_ = *{*2, 3*}*. Here, *G*_*k*_ would include genotypes *g* in *{*0, 1*}*^5^ with the following conditions:

- *g*_2_ = 1 if and only if 2 *∈ l*_*k*_
- *g*_3_ = 1 if and only if 3 *∈ l*_*k*_
- For any other position *i, g*_*i*_ = 0 if *i ∈/ l*_*k*_

Thus, the valid genotypes in *G*_*k*_ would be *{*01000, 00100, 01100, 00000*}*.

We can then define *G* as the union of all such sets:

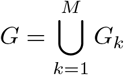

The *epistatic score* for the strategy is given by:

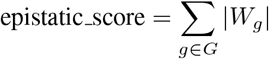

We demonstrate the *epistatic score* of the strategy correlates with its predictive power. This epistatic score quantifies the extent to which large interaction effects between residues are represented in the dataset produced by a specific strategy. PROXICLUST, designed to identify and utilize epistasis clusters within the protein’s binding interface, is expected to perform favorably in terms of epistatic score. Figure 4a presents the epistatic scores for 100 randomly selected assembly strategies, demonstrating a positive correlation between this metric and the R^2^ score of the machine learning models trained using these strategies, with a Spearman correlation coefficient of 0.74. Notably, PROXICLUST achieves superior performance in both R^2^ and epistatic scores compared to the random benchmarks. Results for other datatsets are in SI Figure 7.

**Figure 4:**
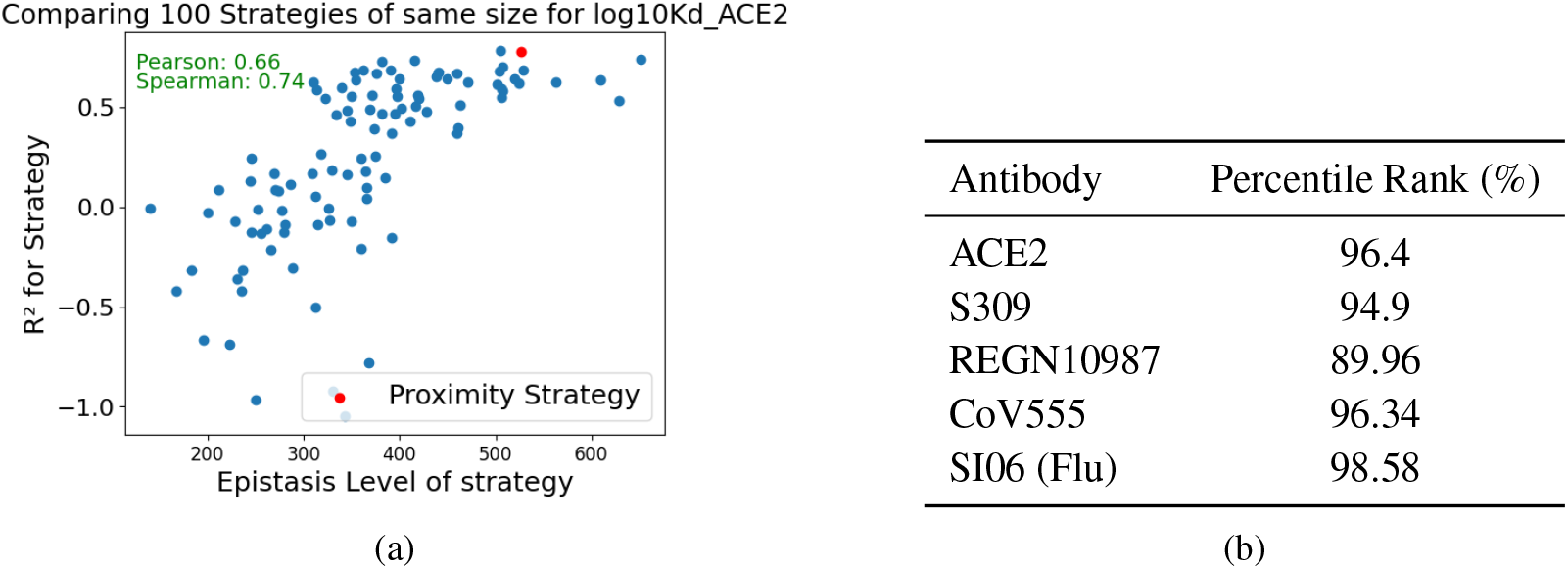
(a) Correlation between model performance (*R*^2^) and epistasis score across 100 random strategies, with PROXICLUST highlighted in red (Spearman: 0.74). (b) PROXICLUST consistently ranks in the top percentiles for epistasis capture across all tested strategies.

Further analysis shown in Figure 4b extends these findings in all five data sets, with PROXICLUST consistently ranking within the top 11% of strategies in terms of epistatic score. The relationship between R^2^ and epistatic score remains robust across these datasets, evidenced by an average Spearman correlation of 0.822 (refer to SI Figure 7).

This comprehensive analysis demonstrates that PROXICLUST’s superior performance arises directly from its systematic ability to identify and capture functionally significant epistatic interactions at the protein binding interface, validating our hypothesis that spatial proximity serves as an effective proxy for mutational interdependence.

## 4 Discussion

This study has demonstrated the potential of using smaller, targeted combinatorial datasets to predict larger, more complex combinatorial datasets with high accuracy at the beginning of a pandemic. PROXICLUST utilizes spatial proximity to infer epistatic interactions, has shown promising results and consistently outperformed random strategies across various datasets.This research demonstrates that just two well-designed combinatorial scans can effectively approximate the entire binding affinity landscape. By strategically leveraging spatial proximity between residues, we focus on mutations with the strongest contributions to binding affinity while avoiding the collection of less significant epistatic effects between residues that are distant in space.

A key implication of our work is its potential application in pandemic preparedness. Early in a pandemic, when new viral variants emerge, our method could guide the strategic design of combinatorial libraries to maximize predictive power. By strategically selecting smaller, experimentally feasible combinatorial sets, researchers could more efficiently predict the effects of larger mutation combinations that might appear in future variants. While existing approaches like the BO-EVO algorithm by Hu et al. (2023) use active learning to guide limited experimental efforts during pandemic response, our method is unique in leveraging a priori structural and biophysical intuition to optimize experimental design before any measurements are taken.

Our findings also raise interesting insights about the fundamental nature of protein-protein interactions. The success of proximity-based clustering in capturing epistatic effects suggests that local structural environment, especially on the binding surface, plays a dominant role in determining binding affinity. This is also demonstrated in several recent publications using local environment to predict protein binding, including the HERMES method which showed that holographic encoding of local atomic environments within 10Å of a focal residue can effectively predict binding affinity and stability effects Visani et al. (2024).

The correlation between epistatic scores and predictive performance provides valuable insights into the mechanistic basis of our method’s success. This relationship suggests that spatial proximity and epistasis are a reliable proxy for functional interactions between residues. While epistatic scores cannot be calculated a priori without experimental data, the consistent correlation between spatial clustering and epistatic effects suggests that our proximity-based approach effectively captures functionally important interactions. Furthermore, this correlation offers a potential metric for evaluating future dataset assembly strategies without requiring extensive experimental validation.

However, several limitations and opportunities for future research should be noted. First, while our method has been successfully validated on datasets containing up to 16 mutations - currently the largest available combinatorial datasets - many real-world scenarios involve substantially larger mutation spaces.Kaku et al. (2024) The correlation between spatial proximity and epistatic effects can become more complex with increasing numbers of mutations, and optimal clustering parameters may need to be adjusted. Nevertheless, the consistent success across our diverse test cases and the strong theoretical foundation linking spatial proximity to epistatic interactions suggest that our approach provides a promising framework for tackling larger mutation spaces.

Second, our method currently relies on structural information about the binding interface. While such information is increasingly available through methods like AlphaFold2 Abramson et al. (2024); Bryant et al. (2022), there are still cases where accurate protein complex structures are unavailable. In particular, antibody-antigen complexes and other adaptive immune system interactions remain challenging to AlphaFold modelYin et al. (2022). Developing alternative data assembling strategies for cases without reliable structural information, perhaps based on sequence conservation or other features, could expand the method’s applicability.

## Code Availability

The code used in this study is made publicly available at https://github.com/mbasse0/ProxiClust

