## Supplementary Information for "Spatial Clustering of Interface Residues Enhances Few-Shot Prediction of Viral Protein Binding"

### A APPENDIX

##### REPRESENTATION OF MUTANTS WITH ONEHOT FORMAT

| Position | 339 | 371 | 373 | 375 | 417 | 440 | 446 | 477 | 478 | 484 | 493 | 496 | 498 | 501 | 505 |
|---|---|---|---|---|---|---|---|---|---|---|---|---|---|---|---|
| Wildtype | G | S | S | S | K | N | G | S | T | E | Q | G | Q | N | Y |
| Mutant | G | S | P | F | K | K | S | N | K | E | Q | G | R | Y | H |
| Onehot representation | 0 | 0 | 1 | 1 | 0 | 1 | 1 | 1 | 1 | 0 | 0 | 0 | 1 | 1 | 1 |

Table 1: Example of onehot representation of one mutant among the $2^{15}$ in the dataset.

#### LOW-THROUGHPUT ACQUISITION METHODS

##### RANDOM ACQUISITION

Random acquisition is the default data acquisition method for most machine learning pipelines Hie et al. (2020), where the training set is a subset of the whole dataset, with an equal probability of selecting any element of the set.

##### DIVERSE ACQUISITION ON ONE-HOT SEQUENCE SPACE

Given a set of labelled mutants $S$, the diverse acquisition scheme samples the unlabelled mutant the furthest from the labelled set at each step, to obtain a new labelled set $S'$ with one additional mutant. More precisely, we studied three possible implementations of that idea:

**Min-linkage** $S' = S \cup \{x\}$ where $x = \arg\max_{y \notin S} \min_{z \in S} H(y, z)$

**Max-linkage** $S' = S \cup \{x\}$ where $x = \arg\max_{y \notin S} \max_{z \in S} H(y, z)$

**Mean-linkage** $S' = S \cup \{x\}$ where $x = \arg\max_{y \notin S} \frac{1}{|S|} \sum_{z \in S} H(y, z)$

Where $H$ is the Hamming distance, $H(x, y) = \sum_{i=1}^{n} \mathbf{1}_{x_i \neq y_i}$ , $x$ and $y$ are two strings of equal length, and $x_i$ and $y_i$ are the $i$-th elements of $x$ and $y$, respectively.

##### DIVERSE ACQUISITION ON PROTEIN LANGUAGE MODEL EMBEDDINGS

The *Diverse acquisition on embeddings* sampling method is the same as above but the distance $H$ is defined as the L2 norm on the embedding space of the protein language model ESM-1v Rives et al. (2019), instead of the sequence space. Such a concept of distance in embedding space is similar to "semanticity" of the protein by Hie et al. in Hie et al. (2021), which serves as a indicator of changes in biophysical properties such as $K_D$.

##### SPARSE ACQUISITION ON ESM EMBEDDINGS

The *Sparse Acquisition* sampling consists in selecting variants in the less dense areas of the ESM embeddings space. A kernel density estimator Pedregosa et al. (2011) is trained on the labeled mutants to obtain a density map of the already explored parts of the 1280-dimensional embedding space. Each new variant added to the training set is chosen to have the embedding with the smallest Kernel Density Estimation value.

##### COMPARING DIVERSE ACQUISITION WITH RANDOM ACQUISITION

Figure 5b is obtained from plot 8. This plot is created by measuring the $R^2$ prediction performance of random forest models trained on a series of 60 training sets of sizes ranging from 10 to 500 samples. The measurements are repeated 112 times for different initializations of the 10 initial samples in the training set, chosen at random. The transparent areas surrounding the $R^2$ curves represent the standard deviation of the scores, computed over the 112 repetitions. To generate Figure 5b, an equally spaced range of 50 R² values are chosen and the corresponding size of training set are obtained on plot 8 by joining the successive points with straight lines junctions. Linear regression analysis, using the scipy linregress package, was performed to calculate the linear fit with a slope of 1.31.

##### UNCONSTRAINED DATA ACQUISITION VIA LOW-THROUGHPUT ACQUISITION METHODS

###### Computing the distribution of R² scores for interface and non-interface strategies

Inter-residue distances are calculated using PDB files that provide the precise three-dimensional positions of every atom in the antibody-antigen binding poses for our datasets. The specific files used are 6M0J Lan et al. (2020) for RBD-ACE2 binding, 8J26 Rahman et al. (2023) for RBD-REGN10987 binding, 7XCK Zhao et al. (2022) for RBD-S309 binding, 7KMG Jones et al. (2021) for RBD-CoV555 binding, 7C01 Shi et al. (2020) for RBD-CB6 binding, 5UGY Whittle et al. (2011) for SI06-CH65. Positions of the residues are then approximated by the coordinates of each $C_\alpha$ atom.

The boxplots on the left of the figures in 3 are made from an exhaustive scan of all the assembly strategies of interface residues into two combinatorial scans. The number of loci contained in the combinatorial scans was set to $L = 6$ for the Covid datasets and $L = 8$ for the Influenza dataset to account for higher level epistasis effects. The size of the interface is set to 10 for covid datasets and 12 for the Influenza dataset. Thus, the boxplots on the left in subfigures of Figure 3 summarize the distribution of $\binom{\binom{10}{6}}{2} = 21945$ assembly strategies. Note that PROXICLUST is one of those 21945 strategies since it also uses interface residues. Similarly, the left boxplot on 3d summarizes the distribution of $\binom{\binom{12}{8}}{2} = 122265$ assembly strategies.

Boxplots on the right of subplots in Figure 3 result from a sampling of a part of all assembly strategies. Indeed, results for the exhaustive scan of either $\binom{\binom{15}{6}}{2}$ or $\binom{\binom{16}{8}}{2}$ assembly strategies was too computationally heavy. $10^5$ strategies are sampled which account for $0.8\%$ of the total number of assembly strategies for the SARS-CoV-2 datasets and $0.12\%$ of the total number of strategies for the Influenza dataset.

#### RESULTS OF LOW-THROUGHPUT ACQUISITION

##### UNCONSTRAINED DATA ACQUISITION VIA LOW-THROUGHPUT ACQUISITION METHODS

We aimed to establish how effectively we could predict larger datasets using any smaller subsets and quantify the accuracy of these predictions in terms of the $R^2$ score. Therefore we evaluated various sampling methods to determine the most effective strategy for constructing the training set. In Figure 5a, we see that only the Diverse data acquisition (see methods) consistently outperforms Random Acquisition across all tested training set sizes. Figure 5b shows Diverse Acquisition uses $31\%$ less training samples compared to the Random Acquisition to achieve same R² scores.

Finally, we determined the number of data points required to achieve a satisfactory $R^2$ score using this strategy. Figure 5d shows the training set sizes necessary to attain high $R^2$ values for combinatorial datasets of various sizes. For a dataset with L loci, which contains $2^L$ samples, the number of points needed to effectively predict the entire dataset grows exponentially. This exponential increase highlights a critical point in protein function prediction: the trade-off between the size of the training set and the model's performance, compounded by the high costs of acquiring experimental data. Notably, while these results are illustrated using RBD-ACE2 data, this trend holds consistently across all datasets we evaluated.

Figure 5d demonstrates that employing unconstrained data acquisition methods, such as Random Acquisition and Diverse Acquisition, is feasible only for combinatorial datasets with a limited number of loci. Given that VoCs sometimes exhibits a lot more mutations than this, and conducting combinatorial experiments on $2^{20}$ combinations or generating random mutations for model training is not experimentally viable. Thus, predicting much larger combinatorial datasets requires high-throughput experimental strategies.

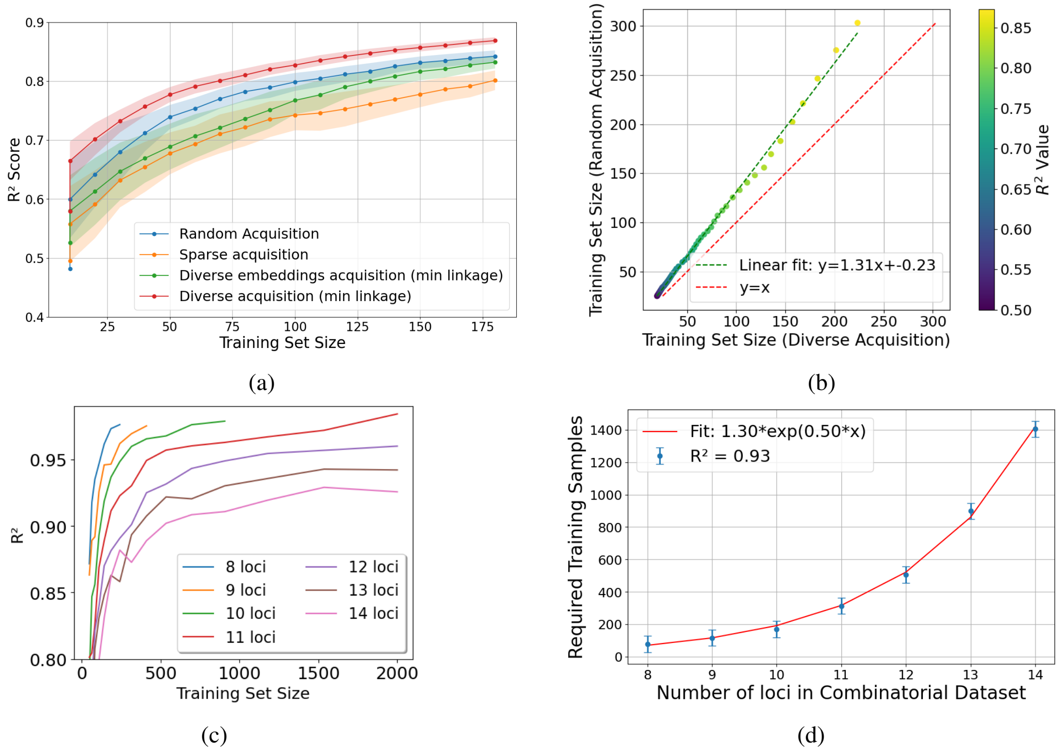


Figure 5: Data acquisition methods comparison (RBD-ACE2 binding affinity prediction). (a) $R^2$ score for varying training set size, compared between four acquisition methods. Only Diverse beats Random. Results are averaged across 112 runs where the initial 10 samples in the training set are initialized differently for each run. (b) Training set sizes required to achieve specific $R^2$ values for Random versus Diverse acquisition method Diverse acquisition saves around 31% of data to achieve similar performance than Random acquisition. (c) $R^2$ scores achieved with random forest models across different training set sizes for datasets of varying complexity (number of loci). (d) Exponential relationship between dataset complexity and required training set size, demonstrating that achieving predictive performance requires exponentially more training data as the number of loci increases.

#### Additional Tables and Figures

| Dataset | Median $R^2$ of all strategies scan | Median $R^2$ of interface strategies scan | $R^2$ score of ProxiClust | Quantile of ProxiClust's $R^2$ score |
|---|---|---|---|---|
| ACE2 | 0.27 | 0.64 | 0.78 | 0.995 |
| REGN10987 | -0.04 | 0.77 | 0.80 | 0.994 |
| CoV555 | -0.05 | 0.68 | 0.73 | 0.897 |
| S309 | -0.53 | 0.41 | 0.73 | 0.986 |
| SI06 | 0.05 | 0.5 | 0.86 | 0.9997 |

Table 2: Results of ProxiClust for 5 different antigen-antibody complexes

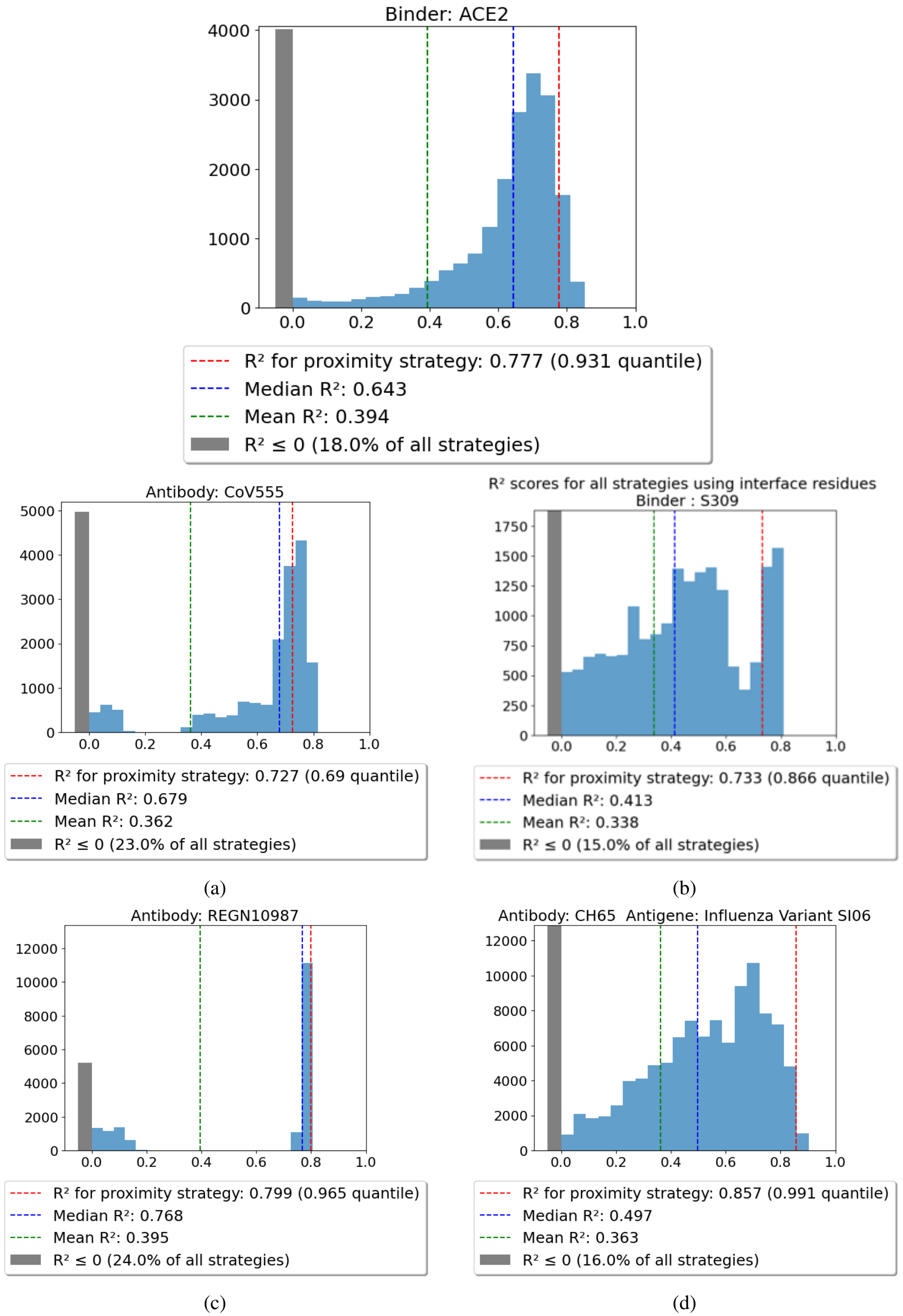


Figure 6: Histograms of $R^2$ scores for models trained from strategies restrained to the antibody-antigen interface. Red line shows that the $R^2$ score achieved by PROXICLUST beats both mean and median values of $R^2$ for strategies sampled at the interface.

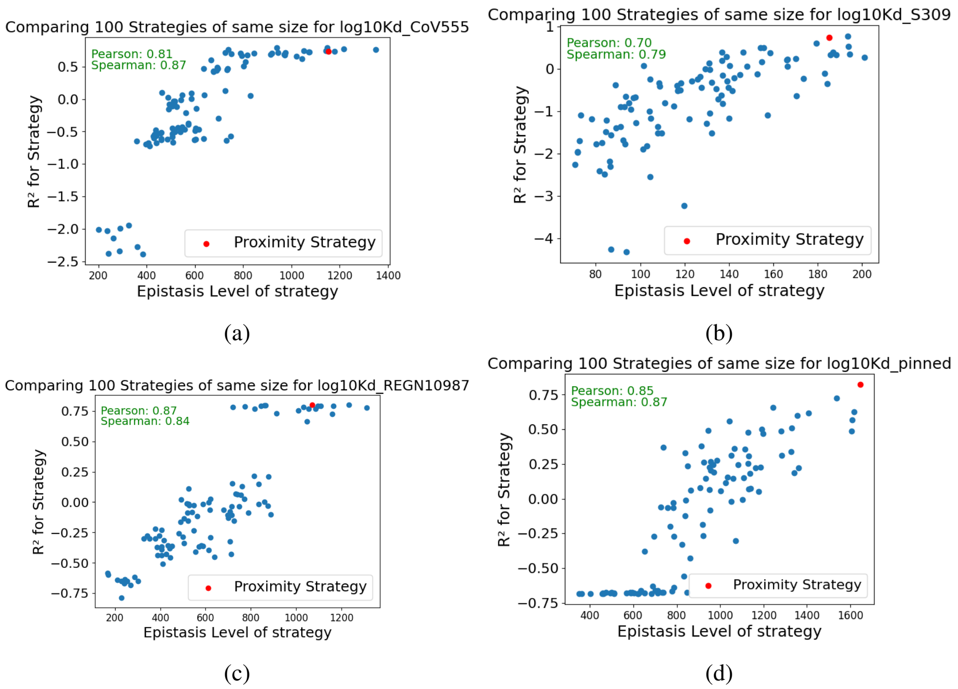


Figure 7: Correlation between model performance and epistasis score across different antibody-antigen binding systems. Each panel shows 100 random mutation selection strategies (blue dots) compared to the ProxiClust strategy (red dot) for: (a) RBD-CoV555, (b) RBD-S309, (c) RBD-REGN10987, and (d) RBD-ACE2 binding. The consistently high Pearson and Spearman correlation coefficients (0.70-0.87) demonstrate that strategies capturing more significant epistatic interactions yield better predictive performance. ProxiClust consistently ranks in the top percentiles for both metrics across all tested systems, validating that spatially-informed clustering effectively identifies functionally important mutational interactions

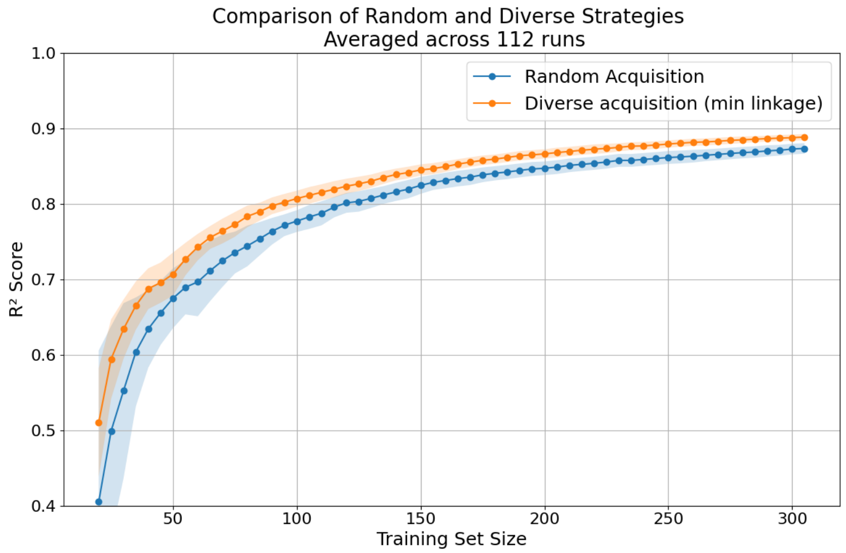


Figure 8: Performance comparison between Random Acquisition and Diverse Acquisition (min linkage) data sampling strategies for RBD-ACE2 binding affinity prediction.
